# kSHREC ‘Delta’ reflects the shape of kinetochore rather than intrakinetochore tension

**DOI:** 10.1101/811075

**Authors:** Fioranna Renda, Valentin Magidson, Irina Tikhonenko, Christopher Miles, Alex Mogilner, Alexey Khodjakov

## Abstract

Distance between fluorescent spots formed by various kinetochore proteins (‘Delta’) is proposed to reflect the level of intrakinetochore tension (IKT). However, larger-scale changes in the kinetochore architecture may also affect Delta. To test this possibility, we measure Delta in long kinetochores of Indian muntjac (IM) whose shape, size, and orientation are discernable in conventional light microscopy. We find that architecture of IM kinetochores and the value of Delta change minimally when microtubule-mediated forces are suppressed by Taxol. In contrast, large decreases of Delta observed in Taxol-treated human cells coincide with prominent changes in length and shape of the kinetochore. We also find that inner and outer kinetochore proteins intermix within a common spatial compartment instead of forming separate thin layers. These observations, supported by computational modelling, suggest that changes in Delta reflect changes in the kinetochore shape rather than the level of IKT.

## Introduction

Segregation of chromosomes during cell division (mitosis) depends upon ‘kinetochores’, macromolecular assemblies located at the centromere of each chromosome. Kinetochores perform two principal functions: they generate the force that propels chromosomes and produce a checkpoint signal that delays progression through mitosis until all chromosomes attach to spindle microtubules. Molecular composition of the kinetochore is complex, comprising over a hundred of various proteins [1]. Further, as the cell progresses through mitosis, molecular composition of the kinetochore and its architecture change in response to various types of interactions with spindle microtubules. These adaptive changes in size and shape of the kinetochores ensure that microtubule attachments form rapidly yet with a low number of errors [2, 3]. Thus, revealing mechanisms that govern kinetochore architecture is of a significant interest.

Due to its small size in most mammalian cells (∼300 nm), the shape of the kinetochore or the distribution of its components cannot be directly delineated in conventional light microscopy (LM). A popular approach to overcoming this limitation is based on measuring the ‘Delta’, a distance between centroids of fluorescent spots formed by various proteins, visualized in different colors within the same kinetochore. Measurements of Delta lay the foundation of a nanometer-scale map that attributes various proteins to thin layers orthogonal to the inner-outer (from the centromere towards attached microtubules) axis of the kinetochore [4, 5]. Delta between the proteins at the base of the kinetochore (CenpA) and those in the microtubule binding domain (Hec1) has been shown to decrease ∼30% when microtubule dynamics are suppressed by Taxol [4, 6]. The decrease was interpreted as a manifestation of changes in the physical separation between the inner and outer layers, which in turn led to the concept of ‘intra-kinetochore tension’ and the notion that stretching the kinetochore is necessary for the satisfaction of the spindle assembly checkpoint (SAC) [7]. This attractive hypothesis; however, remains debatable for two main reasons. First, there exists a high degree of variability in Delta measurements conducted via different techniques and in different laboratories. While some report that Delta (CenpA-Hec1) decreases by ∼30 nm when human cells are treated with Taxol [4], others observe a lesser decrease [8] or no statistically significant change in separation of the same kinetochore proteins under similar experimental conditions [9, 10]. These discrepancies likely arise from the alternative measuring techniques, particularly various approaches to compensating chromatic aberration, inevitable in LM [4, 5, 8, 10, 11]. Second, interpretation of Delta as a metric for physical distances between molecules is obfuscated by the malleable shape of the kinetochore. While literal interpretation of Delta is proven for ‘single-molecule high-resolution colocalization’ (SHREC) analyses of individual molecules [12, 13], applicability of this approach to the kinetochore (SHREC on kinetochores, kSHREC [4]) was originally justified by the layered-disc appearance of kinetochores in Electron Microscopy (EM) [14]. More recent observations of significant alterations in the size and shape of the kinetochore in response to various types of interactions with microtubules [2, 3, 15, 16] challenge this justification [8]. The ongoing debate on the importance of intrakinetochore tension (IKT) prompted us to evaluate relative contributions of distances between kinetochore layers vs. changes in shape and orientation of the kinetochores on the centromere towards the value of Delta. To this end, we applied kSHREC analysis to the compound kinetochores of Indian muntjac (IM) that formed by a lateral fusion of typical mammalian centromeres during evolution of this species. IM kinetochores comprise the conventional thin trilaminar plate (75 nm); however, the length of the plate exceeds 1.5 μm instead of ∼0.3 μm observed in most mammalian cells. The increased length makes the shape and orientation of the kinetochore discernable in LM. Comparative kSHREC analyses in IM vs. human cells suggest that the decrease of Delta observed in Taxol-treated human cells primarily reflects changes in the length of curved kinetochore plates rather that the level of IKT. This prompts re-evaluation of the role ascribed to IKT in the control of mitotic progression.

## Results

### Advantages of Indian muntjac kinetochores for kSHREC analysis

The low number of chromosomes in Indian muntjac (IM) arose from the tandem fusion of numerous normally-sized chromosomes during evolution of this species [17]. As a result, each IM kinetochore comprises tandem repeats of the conventional mammalian kinetochores [18]. While typical mammalian kinetochores appear as small spots in conventional fluorescence LM (Figure 1A,A’), IM kinetochores form thin lines with the length greater than 1 µm (Figure 1B,B’). Molecular composition of IM kinetochores is similar to that of conventional kinetochores [19] and major kinetochore components can be visualized in both via expression of fluorescently-labeled proteins or antibody staining with similar efficiency (Figure 1A,B). At the EM level, IM kinetochores display the typical trilaminar plate with the widths of the layers similar to that observed in other mammalian cells [14, 20, 21]. However, IM plates are several fold longer than the 250-300 nm plates observed in human cells (Figure 1C,D).

**Figure 1.**
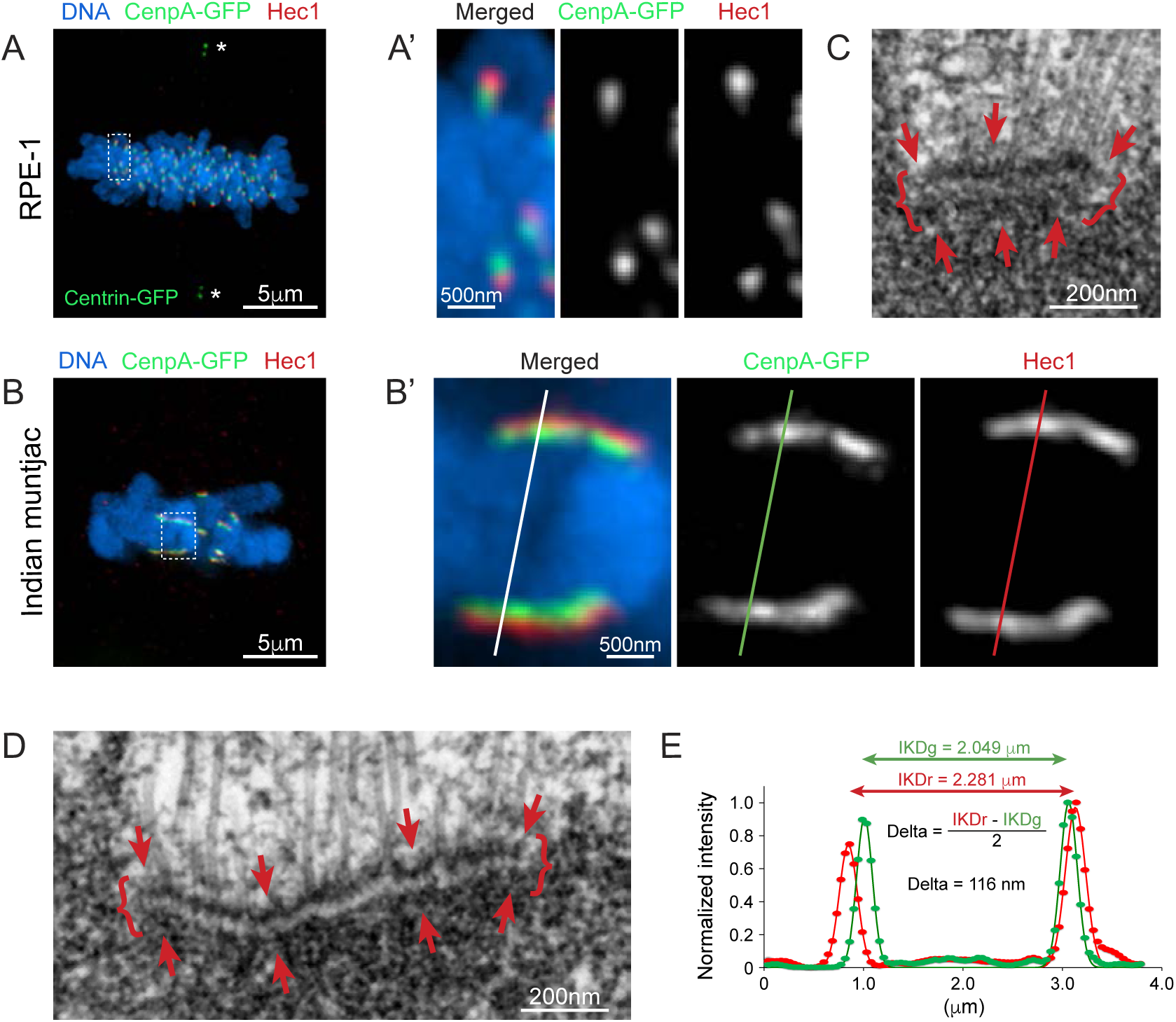
Kinetochore architecture in human and Indian muntjac cells and the approach to Delta measurements. **(A)** Human (RPE1) cell with inner and outer kinetochore domains labeled via expression of CenpA-GFP (green) and immunostaining for Ndc80/Hec1 (red). Paraformaldehyde (PFA) fixation. DNA is counterstained with Hoechst 33342 (blue). Maximum-intensity projection through the entire cell. Centrioles (asterisks) are labeled via expression of Centrin-1-GFP. (**A’)** Higher-magnification of sister kinetochores boxed in A. Both inner (CenpA) and outer (Hec1) kinetochore proteins appear as small spots. (**B)** Indian muntjac (IM) cell with inner and outer kinetochore domains labeled as in A. PFA fixation. (**B’)** Higher-magnification of sister kinetochores boxed in B. Inner and outer kinetochore proteins form thin layers with discernable orientation. Lines denote positions of scan profiles presented in E. (**C)** A 70-nm EM section through a kinetochore in metaphase RPE1 cell. Arrows denote the trilaminar plate comprising ∼25 nm thin electron dense inner and outer layers (arrows) separated by a ∼25-nm thin translucent middle layer. The plate is ∼300 nm long (curly brackets). (**D)** A 70-nm EM section through a kinetochore in IM metaphase. Arrows and curly brackets as in panel C. The plate is ∼1000-nm long and ∼75 nm wide. (E) Line profiles across co-planar sister kinetochores in IM scanned orthogonally to the plates (position of the scan line is marked in B’). Markers are pixel intensities, lines are Gaussian fits. Inter-kinetochore distances IKDg and IKDr are the distances between maxima of the two green and two red peaks correspondingly. Delta is determined as half of the difference between IKDr and IKDg.

Because orientation of sister kinetochores is discernable in IM, intensity profiles can be generated by scanning a line orthogonal to the plates (Figure 1B’). Fitting these profiles with Gaussian functions determines positions of the fluorescent peaks with nanometer precision (Figure 1E) and the distance between the green and red peaks within a kinetochore (i.e., Delta) is then calculated as shown in Figure 1E. Pairwise calculation of Delta for sister kinetochores negates chromatic aberration [4]. Further, accuracy of this approach is not affected by large standard deviations in the distribution of experimental values [8] that have been shown to erroneously increase mean values obtained by measurements of Delta independently for each kinetochore [5, 8]. A major limitation of the pairwise Delta measurement in human cells, is that this approach underestimates the value of Delta if sister kinetochores swivel around the centromere so that their outer layers do not face in opposite directions [4, 8]. However, because the plates of IM kinetochores are discernable in LM, this limitation is overcome by selecting sister kinetochores with the precisely opposite orientation. Thus, Delta in IM can be measured with the level of precision not achievable in kSHREC analyses of smaller human kinetochores whose orientation cannot be accurately determined.

### Delta (CenpA-Hec1) changes minimally in IM cells treated with Taxol

We focus our analyses on the separation between the outer kinetochore protein Hec1 and the centromere-specific histone CenpA that forms the base of the kinetochore. To facilitate direct comparison of the results, the antibodies against Hec1 (9G3, Abcam) and CenpA (3-19, Abcam) are the same as in previous Delta measurements in human cells [4, 5, 8]. We find that Delta_CenpA-Hec1_ in untreated IM metaphase (95±14 nm, Figure 2A) is similar to that previously reported in human metaphase (∼90 nm [5, 8]). Brief exposure to Taxol decreases Delta_CenpA-Hec1_ to 87±14 nm in IM (Figure 2A). This decrease is statistically significant (p = 0.0006 in 2-tailed Student’s t-test); however, it is considerably smaller than ∼20 nm decrease in human cells [5, 8, 22].

**Figure 2.**
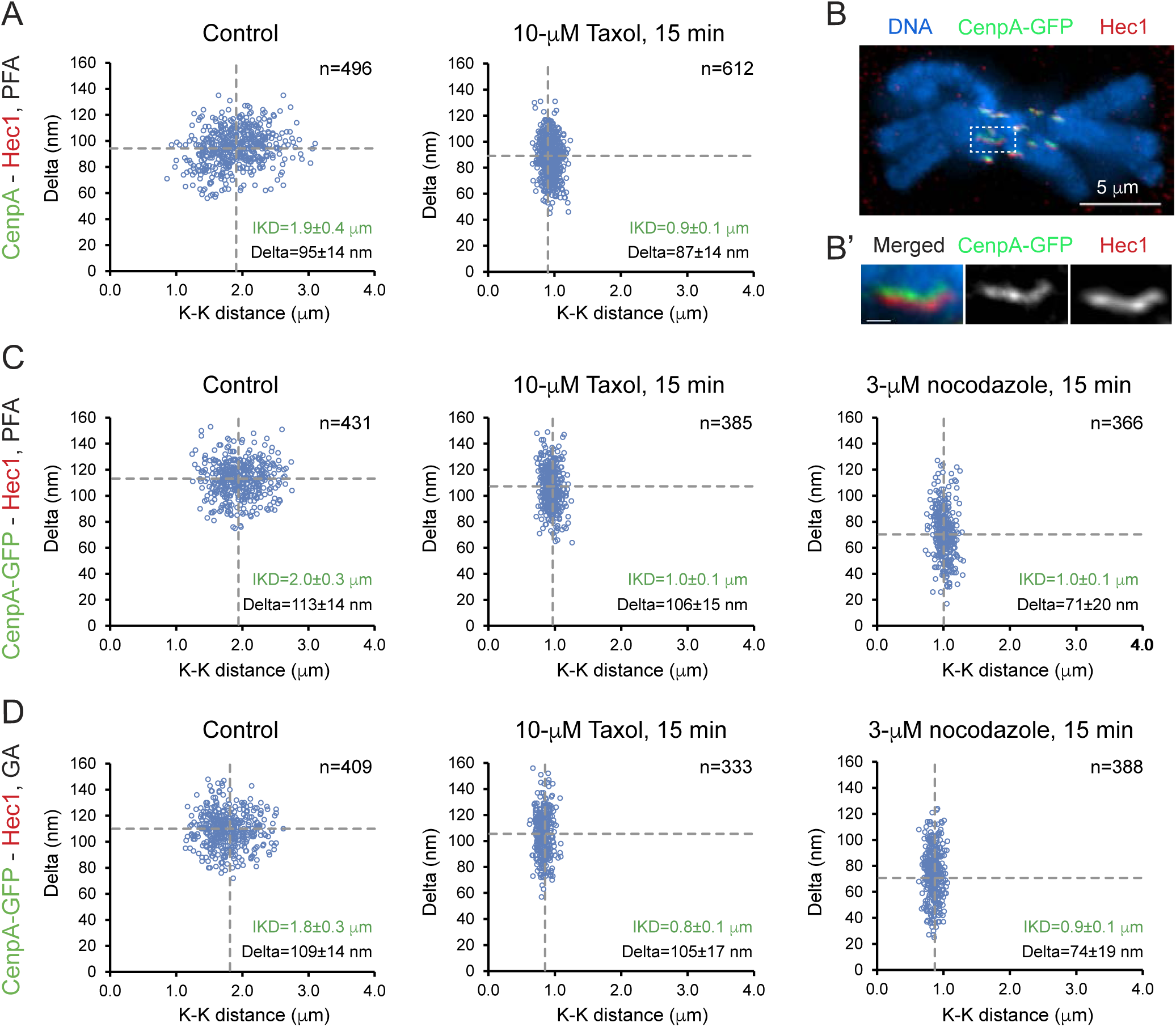
(with 3 supplements). Delta does not decrease significantly in Taxol-treated IM cells. **(A)** Scatterplots of CenpA (green) - Hec1 (red) Delta versus interkinetochore (CenpA-CenpA) distance for untreated metaphase and Taxol-treated IM cells. PFA fixation, followed by methanol. Dashed lines denote mean. Mean ± standard deviation (SD) also shown numerically. (**B)** IM cell with inner and outer kinetochore domains labeled via expression of CenpA-GFP (green) and immunostaining for Ndc80/Hec1 (red). Glutaraldehyde (GA) fixation. Notice that the intensity of staining is similar to Figure 1B although chromosome arms appear slightly more condensed. (**B’)** Higher-magnification of sister kinetochores boxed in B. Scale bar = 500 nm. (**C)** Scatterplots of CenpA-GFP (green) - Hec1 (red) Delta versus interkinetochore (CenpA-GFP – CenpA-GFP) distance for untreated, Taxol-, and nocodazole-treated IM cells. PFA fixation. Mean ± standard deviation (SD) shown. **(D)** As in C but cells are fixed with GA.

Immunolocalization of endogenous CenpA requires partial denaturation of the sample because this protein is buried inside the DNA and thus is not accessible to the antibodies when ultrastructure of the kinetochore is intact. Consequentially, under conditions required for immunolocalization of CenpA, neither microtubules nor the trilaminar appearance of kinetochore plates are preserved as evidenced in EM [23, 24]. To minimize potential effects of sample denaturation we analyze Delta in IM cells that constitutively express CenpA-GFP. Fluorescence of this protein is preserved when cells are fixed with 3.2% PFA (Figure 1B) or 1% GA (Figure 2B, B’), which is important as fluorescent spots containing outer-kinetochore proteins are enlarged in human cells fixed with paraformaldehyde (PFA) vs. glutaraldehyde (GA) [8]. This difference indicates differences in preservation of kinetochore morphology after GA vs. PFA fixation.

We find that Delta_CenpA-GFP-Hec1_ in IM cells fixed with PFA (113±14 nm, Figure 2C) or GA (109±14 nm, Figure 2D) is greater than Delta_CenpA-Hec1_ (95±14 nm, Figure 2A). A similar difference in the distance between Hec1 and CenpA vs. CenpA-GFP positions was observed in human cells [8, 22] and the difference was attributed to the increased fluorescence between the sister kinetochores due to overexpression of CenpA [22]. In contrast to human cells, where Taxol decreases Delta_CenpA-GFP-Hec1_ by greater than 20 nm, in IM the value decreases merely 5-7 nm despite a prominent decrease in the distance between sister kinetochores, which indicates cessation of forces that stretch the centromere (Figure 2C,D). Indeed, inter-kinetochore distances in cells treated with Taxol are similar to the distances in cells where microtubules are completely depolymerized (Figure 2C,D). However, in contrast to Taxol, Delta_CenpA-GFP-Hec1_ decreases by ∼20-nm after 15-min exposure of IM cells to 3-µM nocodazole decreases (Figure 2C,D).

Despite minimal changes of Delta, physiological response of IM to Taxol is similar to that in human cells (Figure 2 – Figure supplement 1). In both cell types, 15-min after addition of 10-µM Taxol, density of spindle microtubules increases prominently near the spindle poles and decreases near the equator (Figure 2 – Figure supplements 1,2,3). During prolonged exposure to the drug, both human and IM cells continue to enter mitosis but remain arrested for longer than 20 hours (Figure 2 – Figure supplement 1; Videos 1 and 2). This behavior indicates that IM cells possess a stringent SAC that is not satisfied despite the minimal decrease of Delta. Thus, our observations are inconsistent with the proposed role of IKT in the control of mitotic progression and/or with the notion that Delta is a reliable metrics for IKT.

### ‘Inner’ and ‘outer’ proteins spatially overlap within IM kinetochores

Morphologically, kinetochores appear as ∼75 nm trilaminar plates in GA-fixed plastic-embedded EM preparations (Figure 1C,D). However, immuno-localization analyses are not consistent with the idea that kinetochore proteins form thin layers within the plate. Correlative immuno-LM/immuno-EM analyses suggest that Hec1 molecules are spread in the direction of the attached microtubules for greater than 200 nm in human cells [8]. A unique advantage of IM kinetochore is that the width of the spatial domains occupied by various kinetochore proteins can be directly measured as the Full Width at Half Maximum (FWHM) of the fluorescence peak in line scans orthogonal to the orientation of the plate (Figure 3A,B).

**Figure 3.**
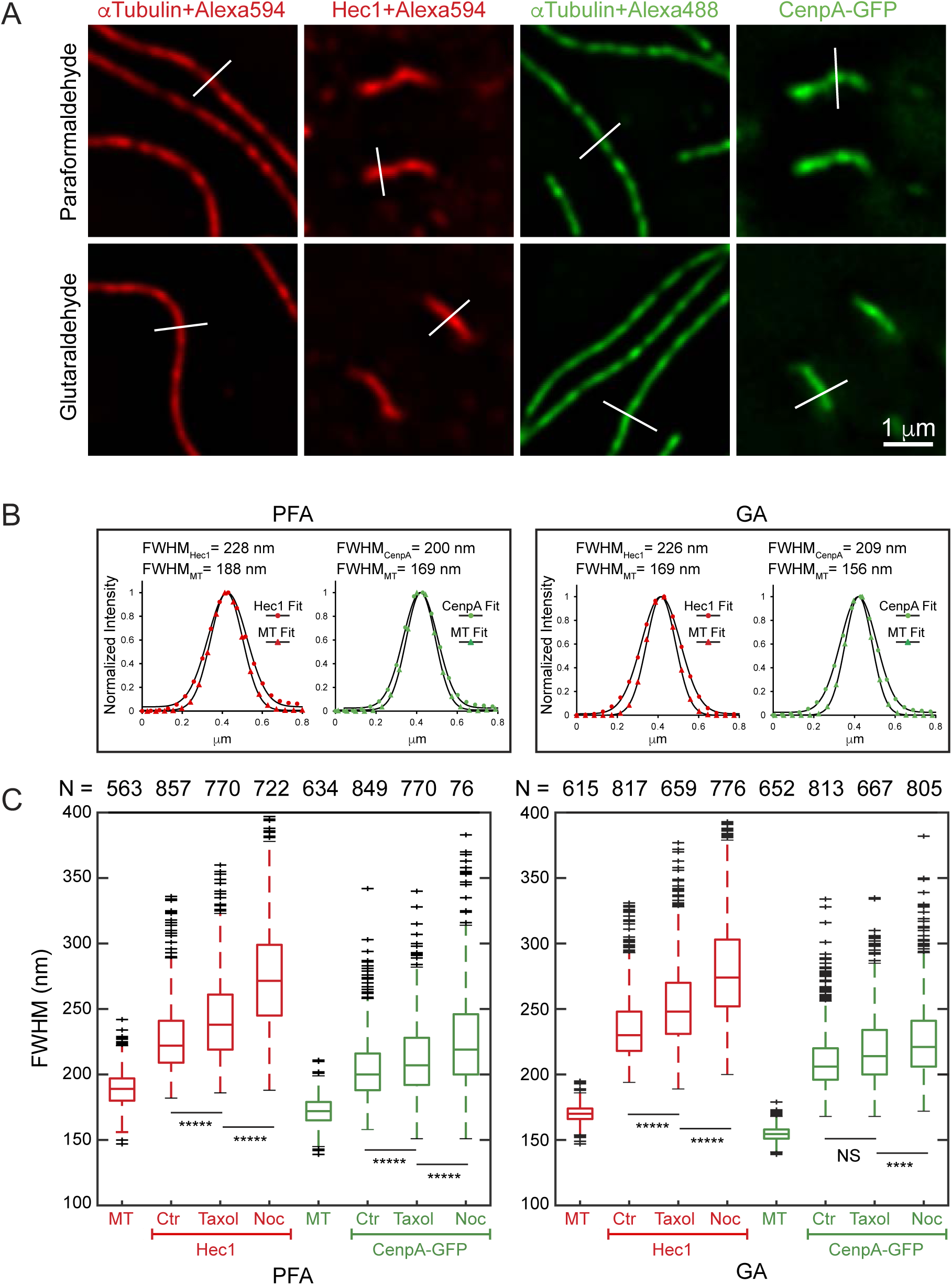
Kinetochore layers in IM cells are wide and their width changes upon Taxol and nocodazole treatments. **(A)** Examples of line-scans (white lines) across individual microtubules or kinetochore plates in cells fixed with Paraformaldehyde (PFA) or Glutaraldehyde (GA). (**B)** Fluorescence profiles corresponding to line-scans shown in A. Markers are pixel intensities, lines are Gaussian fits. Values of the Full Width at Half-Maximum (FWHM) are shown for each profile. Notice that fluorescence peaks of inner- and outer-kinetochore layers are significantly wider than the peaks of individual microtubules. (**C)** Box plots presenting measurements of FWHM for microtubules, inner- (CenpA-GFP), and outer- (Hec1) kinetochore layers in untreated, Taxol- and nocodazole-treated metaphases after PFA (left) and GA (right) fixation. Notice that FWHM of Hec1 layer increases in Taxol- and nocodazole-treated cells. Increase in CenpA-GFP layer is apparent after PFA but not after GA fixation. Student’s t-test p values are less than 10^−4^ (****), 10^−5^ (*****), or greater than 0.5 (NS).

We find that FWHM of the Hec1 distribution within IM kinetochores during metaphase is 233±20 nm after GA fixation and 225±22 nm after PFA fixation (Figure 3C). These values are significantly larger than FWHM of line scans across individual microtubules (170±7 nm after GA and 189±13 nm after PFA, Figure 3C), visualized with the same fluorophore and under identical optical conditions. EM firmly establishes that the diameter of a microtubule in GA-fixed cells is ∼25 nm [25], which implies that FWHM of line scans across a microtubule is determined by the diffraction-limited resolution of the optical system. Thus, the width of Hec1 distribution that exceeds FWHM of diffraction-limited profile by 63 nm, reflects the true physical width of the layer occupied by Hec1 within the kinetochore. Importantly, FWHM values are significantly more variable among Hec1 than among microtubule profiles (Figure 3C). This increased variability supports the notion that the width of microtubule profiles is determined by the optics while the Hec1 profiles reflect natural fluctuations in the organization of the kinetochore plate. We also notice that the width of microtubule profiles in PFA-fixed samples is greater than after GA fixation. This increase likely reflects disintegration of microtubule structure that results in spatial redistribution of tubulin. Indeed, microtubules are not structurally detectable in EM on PFA-fixed samples [25, 26].

Measurements in metaphase cells treated with Taxol demonstrate that abrogation of centromere tension increases FWHM of Hec1 to 251±27 nm after GA and 241±30 nm after PFA fixation. Complete depolymerization of microtubules with high concentration of nocodazole results in a more prominent increase of Hec1 FWHM to 278±35 nm (GA) and 273±39 nm (PFA) (Figure 3C).

CenpA-GFP also localizes within a layer of measurable width. Because resolution is proportional to the wavelength, FWHM is less for a diffraction-limited peak formed by a green vs. red fluorophore. On the microscope used in this study, FWHM of microtubules visualized with a green fluorophore (GFP or Alexa488) is 155±6 nm after GA and 172±11 nm after PFA fixation (Figure 3C). FWHM of CenpA-GFP peaks is significantly wider, measuring 208±18 nm after GA fixation and 202±19 nm after PFA. The width of CenpA-GFP layer increases slightly to 216±24 nm (GA) and 210±26 nm (PFA) in Taxol-treated cells, and further increases to 224±24 nm (GA) and 222±30 nm (PFA) when microtubules are completely depolymerized with nocodazole (Figure 3C).

FWHM measurements demonstrate that both inner (CenpA) and outer (Hec1) kinetochore components reside within layers whose widths are significantly larger than the diffraction limit of resolution and therefore the widths of these layers are measurable in LM. Further, CenpA and Hec1 layers are approximately twofold larger than the distance between the centers of the layers occupied by these proteins (i.e., Delta). These dimensions imply that approximately half of CenpA and Hec1 molecules are spatially intermixed within the same compartment, which in turn means that Delta does not accurately reflect the typical distance between molecules within IM kinetochores.

### CenpA and Hec1 spatially overlap within kinetochores in human cells

Our observation that the width of the layers formed by inner and outer proteins within compound kinetochores of IM exceeds 200 nm raises a question of whether a similar architecture exists in human kinetochores. The underlying assumption in Delta measurements is that kinetochore proteins form negligibly thin layers within an ∼300-nm long plate [4]. A corollary of this assumption is that FWHM of the kinetochore spots in LM should reflect the length of the plate in one direction and be diffraction limited in the orthogonal direction.

To explore whether fluorescently labeled kinetochores in human cells resemble the shape and dimensions assumed in kSHREC analyses, we constructed a computational simulation in which the inner and outer kinetochore layers are modeled as a specified number of ‘molecules’ (points) randomly distributed within a 3-D volume of specified shape. This distribution of molecules is convolved (blurred) with a 3-D Gaussian filter which mimics the point spread function (PSF) of a light microscope. The blurred image is then scaled down to match dimensions of voxels in a typical LM volume recorded on a CCD camera at specified Z-steps. Fitting a Gaussian function to these simulated 3-D images of kinetochores is then used to determine coordinates and FWHM of the peaks (Figure 4A).

**Figure 4 (with 2 supplements).**
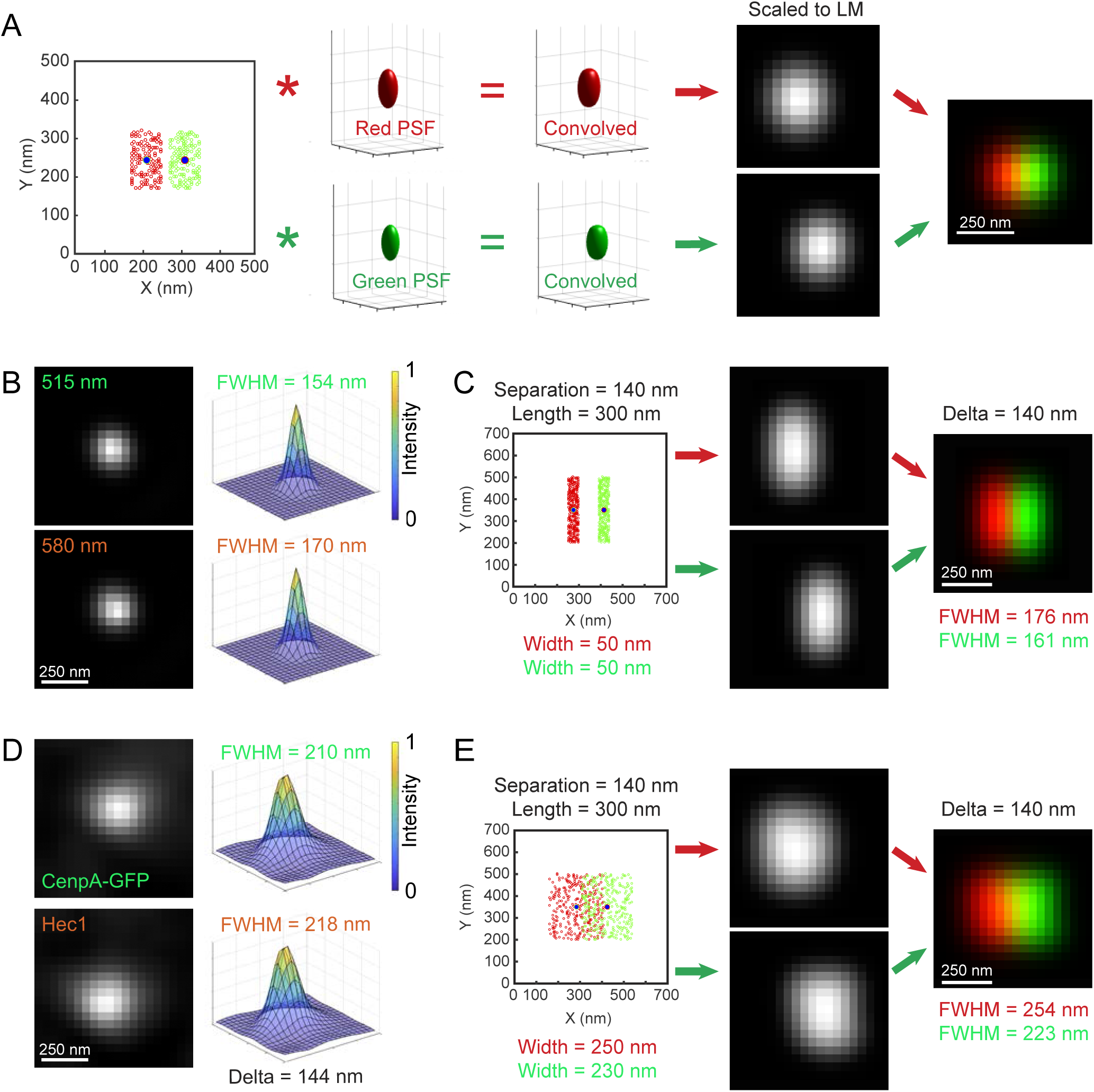
Inner and outer layers of human kinetochores are wide and spatially overlap. **(A)** Layout of computational model used to predict LM appearance of human kinetochores. Randomized distribution of molecules within a specified 3-D shape is convolved with wavelength-specific Gaussian point-spread function (PSF). Convolved volumes are then downscaled to the voxel size typical in conventional LM. (**B)** LM images and intensity profiles of a 100-nm multicolor bead shown in green (515-nm) and red (580-nm) colors. FWHM is reported for the longer axis in XY plane. (**C)** Predicted LM appearance for kinetochores shaped as 300 × 250 nm rectangular prism with 50-nm wide layers separated by 140 nm. Delta and FWHM values are measured in pseudo-LM volume constructed at 40-nm XY pixels and 200-nm Z-steps. (**D)** LM images and intensity profiles of a human kinetochore with typical values of Delta and FWHM (see Figure S2). Notice that both inner (CenpA-GFP) and outer (Hec1) domains are round and significantly larger than a diffraction-limited spot. (**E)** Predicted LM appearance for kinetochores shaped as 300 × 250 nm rectangular prism with widths typical for inner and outer layers observed in IM cells. Centers of the layers are separated by 140 nm. Delta and FWHM values are measured in pseudo-LM volume constructed at 40-nm XY pixels and 200-nm Z-steps.

To establish PSF parameters, we recorded 3-D volumes of multi-color 100-nm fluorescent beads under the same optical conditions as kinetochores (Figure 4B). Measurements of the beads yield 155±27 nm (n = 245) FWHM in the green and 172±31 nm (n = 125) in the red channels. These FWHM values are indistinguishable from FWHM of individual microtubules determined by orthogonal line-scans in GA-fixed samples (Figure 3C) as expected for measurements of diffraction-limited objects on the same microscope. With the experimentally measured PSF parameters, the model predicts that kinetochores comprising inner and outer layers with diffraction-limited widths and separated by Delta previously observed in RPE1 cells [8] should appear as drastically elongated fluorescent spots with minimal overlap between the layers (Figure 4C). In contrast to this prediction, kinetochores appear as more rounded and FWHM of both red and green spots is significantly larger than the diffraction limit, which indicates a noticeable overlap between the inner and outer domains (Figure 4D – Figure supplement 1, also see [8]). This appearance implies that the width of kinetochore layers is similar to the length of the plate. Indeed, computational modelling predicts that for ∼300 long kinetochore plates the width of both inner and outer layers need to be ∼250 nm to match the typical appearance of human kinetochores (Figure 4E). This value is similar to the widths of kinetochore layers measured in the compound IM kinetochores (Figure 3C).

Another feature inconsistent with the notion that proteins form relatively thin layers in human kinetochores is the orientation of the longer axis, particularly for the outer kinetochore components (Hec1, Figure 4 – Figure supplement 1). If these proteins are restricted to a relatively thin but long plate, the longer axis of fluorescent spots formed by these proteins is expected to orient perpendicular to the attached microtubule bundle. In contrast, Hec1 spots are elongated in various directions within the same cell (Figure 4 – Figure supplement 1, Video 3). Variability in the orientation of the longer axes becomes even more prominent in cells treated with Taxol (Figure 4 – Figure supplement 2, Video 4). This change suggests that Taxol alters dimensions and/or shape of the kinetochore, which may affect the value of Delta.

### Kinetochore plates are curved, and the length of plate increases in Taxol treated human cells

Canonical interpretation of Delta as metrics for the distance between layers of molecules assumes that the overall shape and dimensions of the kinetochore plate remain constant. Previous computational analyses suggest that for inflexible layers Delta accurately reflects the distance even if the layers are slanted or jagged [4]. However, for curved layers, the value of Delta is smaller than the physical distance between the layers and the magnitude of this difference is greater for curvatures with smaller radii and/or longer plates (Figure 5A, also see reference [8]). Systematic analysis of the kinetochore plate length and curvature in EM preparations has not been reported, which prompted us to assess these parameters in serial 70-nm plastic sections of untreated (Figure 5 – Figure supplement 1) vs. Taxol-treated (Figure 5 – Figure supplement 2) RPE1 cells.

**Figure 5 (with 2 supplements).**
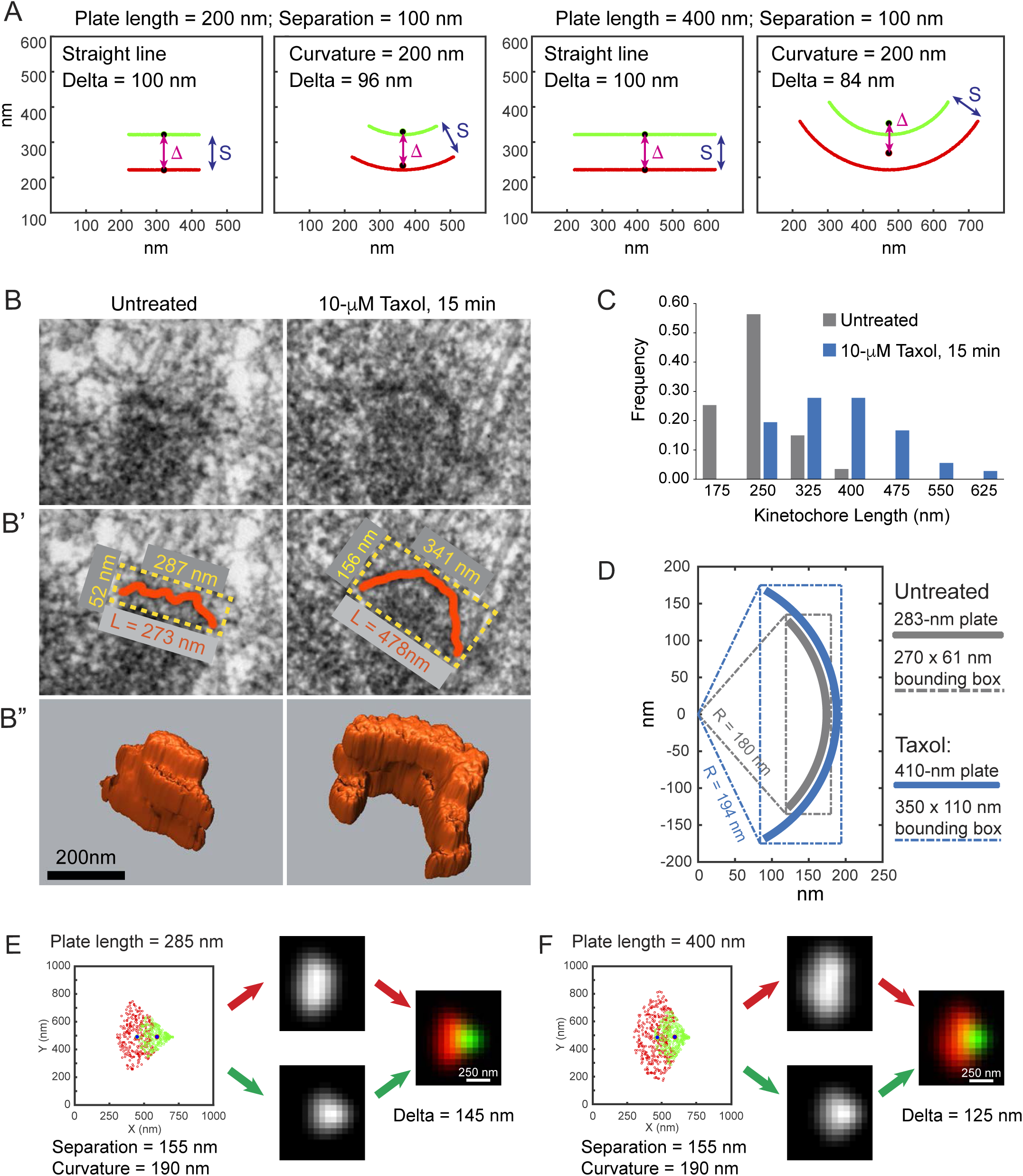
Kinetochore plates are curved and their length but not curvature increases in Taxol-treated human cells. **(A)** Effects of the plate curvature and length on the value of Delta. For parallel straight lines of any length value of Delta matches the distance separating the lines. In contrast, for circular arcs, Delta is smaller than the distance separating the lines and the difference depends on the curvature and length of the arc. (**B)** 70-nm EM sections through typical kinetochores in untreated (left) vs. Taxol-treated RPE1 cells (see Figure 5 – Figure supplements 1,2). **(B’)** Kinetochore plate in Taxol-treated cell is longer and the outer layer fits within a rectangular box of greater length and width than in untreated cell. **(B’’)** Surface-rendered reconstructions from serial sections demonstrate complexity of the outer plate shape in 3-D space. (**C)** Frequency of plates with various lengths in untreated (N = 87 kinetochores) and Taxol-treated (N = 38 kinetochores) cells. Mean values of plate length are 283±54 in control and 410±84 in Taxol-treated cells. (**D)** Dimensions of the bounding box required to encompass the outer plate in untreated (270±47 by 61±18 nm) vs. Taxol-treated cells (350±57 by 110±39 nm) suggest that the outer plate resembles a circular arc with ∼190-nm radius. (**E)** Predicted LM appearance for kinetochores that comprise 250-nm wide layers with length and curvature observed in untreated RPE1 cells. Notice that centers of the green and red layers need to be separated by ∼155 nm to achieve the expected 145-nm Delta. **(F)** Predicted change in the LM appearance and Delta due to increased length of the plate. All other simulation parameters are identical to (E).

Tracing kinetochore plates in GA-fixed conventionally embedded cells (Figure 5B) demonstrates that the length measured in sections oriented parallel to the coverslip surface is 283±54 nm (N = 87) during metaphase but it increases to 410±84 (N = 38) when metaphase cells are treated with Taxol for 15 min (Figure 5C). Both values are significantly greater than FWHM of Hec1 spots measured in LM (Figure 4 – Figure supplements 1,2).

The shape of outer kinetochore plate is highly variable both in untreated and Taxol-treated cells (Figure 5 – Figure supplements 1,2). Undulations, protrusions, and cavitations are prominent in most kinetochores. To estimate a typical curvature of the plate we determined dimensions of the minimal rectangular box that fully encompasses the plate in a single section (Figure 5B’). This approach yields more uniform results than direct fitting of the plate with a circular arc. The latter produces large residual errors due to minute irregularities in the shape of the plate, particularly when the plate is short (our unpublished observation). We find that a typical outer kinetochore plate in untreated cells is encompassed by a 270±47-nm by 61±18-nm rectangle while Taxol-treated kinetochores require a 350±57-nm by 110±39-nm bounding box (Figure 5D). Assuming a perfectly circular arc, these dimensions suggest that typical kinetochore curvatures are ∼180 nm for the untreated and ∼194 nm for Taxol-treated cells (Figure 5D). The difference between these values is not statistically significant.

Thus, EM analyses suggest that a typical kinetochore plate is an arc with the curvature radius of ∼190 nm. Taxol does not significantly change the curvature, but it prominently increases the length of the plate. To more clearly understand how these geometric features of the kinetochore plate influence the results of FWHM analyses, we employed a computational model. The model makes two major predictions. First, to achieve ∼145-nm Delta observed in untreated RPE1 cells, kinetochore layers need to be separated by ∼155 nm (Figure 5E). Second, elongation of the plate with fixed curvature leads to a decrease in Delta from 145 nm to 125 nm (Figure 5F). The model also predicts LM appearance of kinetochore spots that resembles appearance of real kinetochores in untreated and Taxol-treated RPE1 cells (Figure 5E,F, Figure 4 – Figure supplement 1,2).

## Discussion

Pioneering time-lapse recordings of live cells with kinetochores labeled in two different colors revealed that the distance between fluorescent spots corresponding to the microtubule-binding domain and the base of kinetochore was not constant. Specifically, the spots appeared to be closer together when microtubule dynamics were suppressed with Taxol, a treatment known to reduce the force that stretches centromeres of bioriented chromosomes [6, 27]. These observations gave birth to the concept of IKT [7]. Subsequent pairwise measurements of distances between the spots formed by various kinetochore proteins (i.e., Delta) laid the foundation of a linear map that ascribed various proteins to specific layers within the kinetochore and interpreted Delta as the distance that separates these layers [4]. Essential for this interpretation of Delta was the assumption that kinetochores are flat thin plates whose shape and dimensions remain constant upon Taxol treatment. Here we test this assumption and find that a typical kinetochore plate in human cells is curved, which implies that Delta underestimates the distance between the layers. We also find that the length of curved kinetochore plates increases in Taxol-treated cells. This type of architectural reorganization inevitably decreases the value of Delta even if the distance separating inner and outer kinetochore layers remains constant. Thus, our data demonstrate that Delta is not directly proportional to IKT.

Our observation of a significant plate curvature is consistent with early EM descriptions of kinetochores as crescents with the convex side facing away from the centromere [reviewed in 28]. Indeed, kinetochore plates in human cells cannot be truly flat because they protrude from the surface of a cylindrical centromere, whose relaxed radius is less than 400 nm. A force pulling kinetochores outwards is expected to increase the plate’s curvature and our EM results suggest that the typical radius of kinetochore plates visualized via conventional EM is less than 200 nm (Figure 5). At this curvature Delta would underestimate the distance between negligibly thin layers by ∼20% (Fig5A). However, our data suggest that kinetochore layers are not negligibly thin, and the widths of the layers are not constant – they increase under conditions that decrease Delta (Figure 3, also see [8]). These complex architectural changes further complicate quantitative interpretation of Delta.

FWHM measurements of Hec1 and CenpA spots in human cells also suggest a significant spatial overlap between the inner and outer proteins. Our data demonstrate that human kinetochores are much larger than diffraction-limited spots in all directions, i.e., the width of the kinetochore is as large as its length. This observation is supported by computational simulations with experimentally-determined parameters of PSF. These simulations predict that fluorescent spots produced by protein layers constrained within flat thin discs do not resemble appearance and dimensions of kinetochore spots observed in human cells (Figure 5E, Figure 4 Figure supplement 1, 2). Thus, ‘inner’ and ‘outer’ proteins are not separated by a typical distance represented by Delta but intermixed within a common spatial compartment.

Consistent with the notion that large-scale architectural reorganizations contribute significantly towards the value of Delta in human cells, we observe that Delta remains virtually constant in Taxol-treated IM cells. Due to the increased length of the kinetochore as well as a larger diameter of centromere in IM, kinetochore plates in these cells are relatively flat (Figure 2 Figure supplement 2) and their shape is largely preserved in Taxol-treated cells (Figure 2 – Figure supplement 3). These features ensure that Delta values determined in line-scans orthogonal to the plate accurately reflect the distance between the inner and outer kinetochore domains. Thus, minimal changes in Delta values upon suppression of microtubule forces by Taxol in IM cells imply one of the two possibilities. First, mechanic properties of IM kinetochores may fundamentally differ from those in humans. Second, the large decrease in Delta observed in human cells primarily reflects a change in overall shape/size of the kinetochore and not the distance between the layers of molecules. Similar physiological response of IM and human cells to Taxol, similar molecular composition of kinetochores [19], similar morphology of kinetochore plates in both species (Figure 1), and similar values of Delta in untreated cells (cf. Figure 2 and [4]) render the first possibility unlikely. Therefore, Delta in human cells appears to reflect primarily larger scale deformations of malleable outer plate rather than elastic stretching within the plate (i.e., IKT). Our interpretation gains additional support from observations of different Delta values in sister kinetochores [8, 27, 29]. It is impossible for linearly connected elastic elements to simultaneously experience different levels of tension. In contrast, changes in the shape of kinetochore plate are expected to be independent for sister kinetochores and prominent differences in the shape of sister kinetochores is directly observed in EM (Figure 5 – Figure supplements 1,2).

In recent years, kSHREC analyses have led to contradictory conclusions regarding the nature of IKT as well as its role in mitotic progression. Some studies attribute 100% of changes in Delta to changes in the distance between various kinetochore proteins or conformational changes within a specific molecule [4]. Others conclude that Delta in part [8] or exclusively [10] reflects swiveling (angular reorientation) of kinetochores on the centromere instead of reporting correct intrakinetochore distances. These discrepancies are exaggerated by large differences in absolute values of Delta for the same pair of proteins that vary as much as 40% in different reports [4, 5, 8, 10]. In part, this lack of consensus may be due to different approaches to Delta measurements employed in different laboratories and there is a thoughtful debate on the conditions necessary for accurate localization of fluorescent spots in cells [5, 11]. However, a larger issue is whether the size of kinetochore is sufficiently small and whether the shape of this organelle is sufficiently invariant to allow unequivocal interpretation of Delta as the distance between various molecules. This issue has not been considered since the introduction of kSHREC approach a decade ago [4]. Our data presented here suggest that human kinetochores are too large, complexly shaped, and too flexible to be amenable to Delta measurements. While center-of-mass positions for various molecules can be determined with sub 10-nm precision, interpretation of Delta as the distance between molecules or the length of a molecule within the kinetochore [5] is valid only if the molecules form a parallel array within a rigid disc of constant dimensions. Thus, currently there is no proof that kinetochores are amenable to SHREC-based analysis. This urges caution in literal interpretation of Delta as metrics for IKT or distance between subdomains within the kinetochore.

## Materials and Methods

### Cell culture and drug treatments

Immortalized hTERT-RPE1 constitutively expressing CenpA-GFP and Centrin-1-GFP [30] and hTERT-Indian Muntjac cells ([19], kind gift of Dr. Helder Maiato, Institute of Molecular and Cellular Biology, Porto, Portugal) were cultured in DMEM (Gibco, Life Technologies) supplemented with 10% FBS (Sigma-Aldrich F0926) at 37°C and 5% CO_2_. Taxol (Paclitaxel T7402-5MG; Sigma-Aldrich) and nocodazole (Calbiochem, #487928) were added ∼ 15 min prior to fixation to final concentrations of 10-μM and 3-μM respectively.

### Time-lapse recordings

Cells were seeded in 12.5 cm^2^ Tissue Culture Flasks and grown in DMEM (Gibco, Life Technologies) supplemented with 10% FBS (Sigma-Aldrich F0926) and 25 mM HEPES (Gibco by Life Technologies REF 15630-080) at 37°C and 5% CO_2_ for 24 h. Taxol was added in conditioned culture media to 5 μM final concentration, 30 minutes prior to filming. Cells were imaged on Nikon TS100 microscopes equipped with a 10X objective lens (Nikon) at 37°C for 48-72 h. Phase-contrast or Differential Interference Contrast images were captured at 2 min intervals on a Monochrome Spot IR camera. The microscope and light source were controlled by Spot 5.3 Advanced software (Diagnostic Instruments, Inc.).

### Immunofluorescence

Cells were concurrently permeabilized and fixed in PEM buffer (100 mM Pipes, pH 6.9, 2.5 mM EGTA, and 5 mM MgCl2, pH 6.9) supplemented with 1% Triton X-100 and 1% glutaraldehyde (G5882; Sigma-Aldrich) or 1% Triton X-100 and 3.2% paraformaldehyde (EM grade; EMS) for 10 min. Microtubules were visualized with DM1α monoclonal anti-α-tubulin antibody at 1:100 (T9026; Sigma-Aldrich) followed by a secondary antibody conjugated with Alexa Fluor 594 (A-11032; Thermo Fisher Scientific) or Alexa Fluor 488 (A-11029, Thermo Fisher Scientific). Outer kinetochore was stained with a monoclonal 9G3/Hec1 antibody (Abcam ab3613) at 1:1000 followed by a secondary antibody conjugated with Alexa Fluor 594 (A-21135; Thermo Fisher Scientific). In the experiments that involved immunostaining for endogenous CenpA, the cells were fixed in paraformaldehyde and postfixed in 100% methanol at –18°C for 15 min. Endogenous CenpA was visualized with a mouse monoclonal antibody (3-19, Abcam; ab13939) at 1:200. Although both 3-19 (CENP-A) and 9G3 (Ndc80/Hec1) antibodies are mouse monoclonal, their isotypes are distinctly different, which allows highly specific co-staining. 3-19 antibody was detected with IgG1 (γ1)-specific secondary antibody conjugated to Alexa Fluor 488 (Thermo Fisher Scientific, A-21121;), and 9G3 was detected with IgG2a (γ2a) specific secondary antibody conjugated to Alexa Fluor 594 (Thermo Fisher Scientific, A-21135;), both at 1:100 dilution.

Immunostained cells were embedded in non-solidifying media containing 90% glycerol, 10% 1M Tris-Cl, pH 8.5, and 1 mg/ml PPDA. Wide-field fluorescence images were obtained on a Nikon TE2000E2 microscope equipped with 100x 1.49 NA PlanApo TIRF lens and LED illuminator (CoolLED PE 4000). Images were captured with an Andor Zyla 4.2 camera at 43-nm X-Y pixels and 200-nm Z steps. The system was controlled by Nikon NIS-Elements Advanced Research software (Nikon instruments). All images were deconvolved with SoftWorRx 5.0 deconvolution software (Applied Precision), and objective lens-specific in house-recorded point-spread functions. Deconvolution method was set to “Conservative”, noise level to “Medium” for kinetochores and “Low” for 100-nm beads, and the process ran for 10 iterations.

### Serial section electron microscopy

IM and RPE1 cells were fixed with 2.5% glutaraldehyde (G5882; Sigma-Aldrich) in PBS, pH7.4 for 30 min, rinsed with PBS (3 × 5 min), and post-fixed with 2% OsO_4_ in dH_2_O for 60 min at 4°C. The coverslips were then rinsed in dH_2_O, treated with 0.25% tannic acid for 20 min, and stained with 2% uranyl acetate for 60 min. Dehydration was achieved by a series of ethanol solutions (30-50-70-80%–96%, 10 min in each solution) followed by acetone (10 min). After dehydration, cells were embedded in Epon 812 and cured for 48 hours at 60°C. Serial 70-nm thin sections were cut with a diamond knife (Diatome) on a Leica Ultracut UCT ultramicrotome and stained with lead citrate. Images were obtained on a JEOL 1400 microscope operated at 80 kV using a side-mounted 4.0 Megapixel XR401 sCMOS AMT camera (Advanced Microscopy Techniques Corp). Higher-magnification images (40K) were collected for individual kinetochores. These high-magnification images were subsequently used to trace kinetochores plates. 3-D volumes occupied by the kinetochores were visualized as isosurface models in Amira 5.3.3 (Visage Imaging).

### Measurement of interkinetochore distances, Delta, and Full Width at Half Maximum (FWHM) in IM cells

Intensity profiles of selected centromeres (CenpA-GFP) with co-planar sister kinetochores (Ndc80/Hec1) were obtained for single-pixel lines ImageJ. Precise position and FWHM of the two Gaussian peaks in each of these profiles were obtained via the Peakfit MATLAB function developed by Dr. O’Haver (https://www.mathworks.com/matlabcentral/fileexchange/23611-peakfit-m). Profiles with fit errors larger than 3% were discarded (less than 3% of all profiles).

Interkinetochore distance (IKD) was calculated by subtracting the distance between the centers of CENP-A or Hec1 peaks. Delta values were calculated by subtracting the distance between the centers of CENP-A peaks from the distance between the centers of Ndc80/Hec1 peaks and dividing the result by 2. In this approach, potential errors due to chromatic aberration are automatically negated by the opposite orientation of sister kinetochores. However, the same (average) value of Delta is assigned to both sister kinetochores.

### Measurement of interkinetochore distances, Delta, and Full Width at Half Maximum (FWHM) in RPE1 cells

Precise positions and dimensions of fluorescent spots in RPE1 cells were obtained with the delta_tool MATLAB function described in reference [8] and available at http://www.jcb.org/cgi/content/full/jcb.201412139/DC1. This function determines x-y coordinate of the kinetochores center of mass via nonlinear fitting (lsqcurvefit MATLAB function) in projections of segmented volumes of kinetochores with a 2D Gaussian function. Z coordinates are estimated separately in x-z projections. Chromatic aberration errors are suppressed by shifting one color channel so that the global centers of mass for the red and green channels coincide. Delta is then calculated directly as the distance between the red and green centroids within the same kinetochore.

### Measurement of the outer plate length and curvature

Length of the kinetochore outer plate was determined by tracing the contours of the plate with the Freehand line tool in ImageJ (National Institutes of Health). To determine curvature of the outer plate, a minimal rectangular box that encompasses the entire plate was drawn with the Rectangle tool of ImageJ. For a perfectly circular segment, curvature radius of an arc constraint within a rectangular box is calculated as R = W^2^/8H+H/2 where W is the length of the rectangle that defines the chord, and H is the height of the rectangle that defines the sagitta of the arc.

### Computational simulation for evaluation LM appearance of kinetochores with various architecture

The simulation is made using a Matlab code, and the following parameters are considered. Kinetochores are modeled as two plates, each consisting of fixed number of molecules uniformly distributed randomly within a 3-D volume of specified dimensions and position relative to the other plate with the possibility of overlap. The volume is rectangular except for curvature in either the XY or XZ planes with assumed rotational symmetry with respect to both the Z and Y axes. Two PSFs for “green” and “red” colors are constructed by convolving a single point with 3-D gaussian filter (implemented via fspecial3 function available in Matlab 2018b or a later version of Matlab). Widths of the PSFs are specified independently in XY plane and along Z direction. Convolved volumes are down-sampled to the XY pixel size and Z steps expected in LM by integrating intensity constrained within the part of convolved volume that corresponds to the specified LM voxel. Relative position of the down-sampled LM array with respect to the full-resolution convolved volume is randomized within 1 LM pixel to mimic random positions of kinetochores with respect to CCD pixels. Pseudo-experimental measurement of Delta and FWHM are conducted as in real experiments but within the simulated volume at LM resolution. Simulation code is available in the supplemental materials.

### Preparation of illustrations

Contrast and brightness of the final LM images were linearly adjusted in Photoshop (CS6) and the figures assembled in Illustrator (CS6; Adobe). Graphs were prepared in MATLAB or Excel and imported into Illustrator as PDFs.

## Acknowledgements

We thank Dr. Helder Maiato (i3S, Porto, Portugal) for the kind gift of IM cells expressing CenpA-GFP. Electron microscopy was performed with the help of the Wadsworth Center’s Electron Microscopy Core Facility. This work was supported by R35 GM130298 to A.K.

## Video legends

**Video 1. Effects of Taxol on RPE1 cells.** Cells that enter mitosis (manifested by the characteristic round morphology) in the presence of 5-μM Taxol remain arrested and eventually die. Phase-contrast microscopy. Time stamp in hours: minutes. Selected time points from this recording are shown in Figure 2 - figure supplement 1C.

**Video 2. Effects of Taxol on IM cells.** Taxol treatment and filming conditions identical to Video 1. Selected time points from this recording are shown in Figure 2 - figure supplement 1C.

**Video 3. Distribution and appearance of kinetochores during metaphase in RPE1 cell.** Numbers denote centromeres with sister kinetochores indexed as ‘a’ and ‘b’. Each kinetochore is labeled in green (CenpA-GFP) and red (Hec1 visualized with Alexa594). Centrosomes (Ca and Cb) are labeled with Centrin1-GFP. Chromosomes are stained with Hoechst 33342 (blue). Scale bar in the bottom-right corner applies to X–Y planes; depth of each plane is denoted in the top-left corner. All 92 kinetochores from this volume are shown at high magnification in Figure 4 - figure supplement 1. Notice that spatial proximity of kinetochore 7a to 8a as well as 17a to 18a preclude accurate Delta and FWHM measurements for centromeres 7, 8, 17, and 18. Glutaraldehyde fixation.

**Video 4. Distribution and appearance of kinetochores in Taxol-treated RPE1 cell.** Taxol added ∼15 min prior to fixation when the cell had assembled a bipolar spindle with congressed chromosomes. Fixation, staining, and imaging conditions identical to Video 3. All 92 kinetochores from this volume are shown at high magnification in Figure 4 - figure supplement 2. Notice that Delta and FWHM measurements were not obtained for centromeres 7-9, 11, 27, 30, 33, and 38-42 due to spatial overlap among their kinetochores. Glutaraldehyde fixation.

## Source-data files

**Figure 2 – source data 1**. Numeric values for data presented in panels A, C, and D. Data for each plot are compiled in a tab within MS Excel file.

**Figure 3 – source data 1**. Numeric values for data presented in panel C. Data for each experimental condition are compiled in a tab within MS Excel file.

**Figure 4 – source data 1**. Computer code used to generate panels A, C, and E. Requires MATLAB 2018b to run.

**Figure 5 – source data 1**. Numeric values for panels C and D. MS Excel format.

**Figure 5 – source data 2**. Computer code used to generate panels E and F. The same code is also used in Figure 4. Requires MATLAB 2018b to run.

## Supplemental Figures

**Figure 2 – figure supplement 1.**
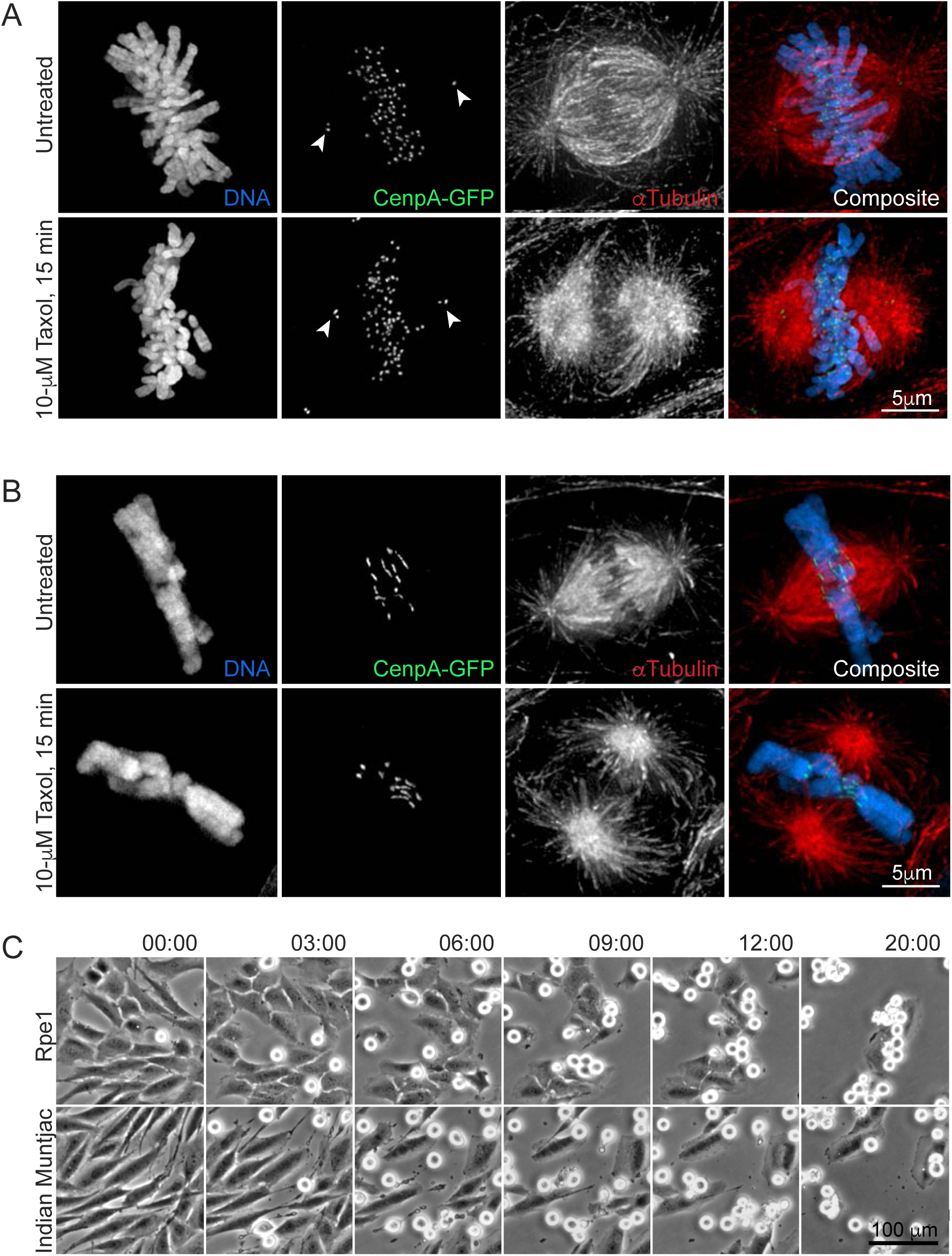
Effects of Taxol in IM and human cells. **(A - B)** Similar changes in mitotic spindle architecture in human Rpe1 (A) and IM (B) cells. Approximately 15 min after addition of Taxol (10 μM final concentration), already assembled mitotic spindles remain bipolar with congressed chromosomes in both human and IM cells. Microtubules density decreases near chromosomes and increases near spindle poles (A) Maximum-intensity projection (MIP) of an untreated (top) and Taxol-treated (bottom) Rpe1 metaphases. (B) MIP of an untreated (top) and Taxol-treated (bottom) IM metaphases. Kinetochores are labeled via expression of CenpA-GFP (green); microtubules are immunostained with a monoclonal antibody (DM1A, Sigma) against α-tubulin (red); chromosomes are stained with DNA dye Hoechst 33342 (blue). Rpe1 cells additionally express Centrin1-GFP that marks positions of centrioles/spindle poles (arrowheads in A). **(C)** Stringent mitotic arrest in Rpe1 (top) and IM (bottom) in the presence of 5-μM Taxol. In both cell types mitosis is arrested for >20 hrs and most cells subsequently die. Selected video frames from a 48-hr long time-lapse recording at 2-min intervals (see Videos 1 and 2 for full recordings).

**Figure 2 – figure supplement 2.**
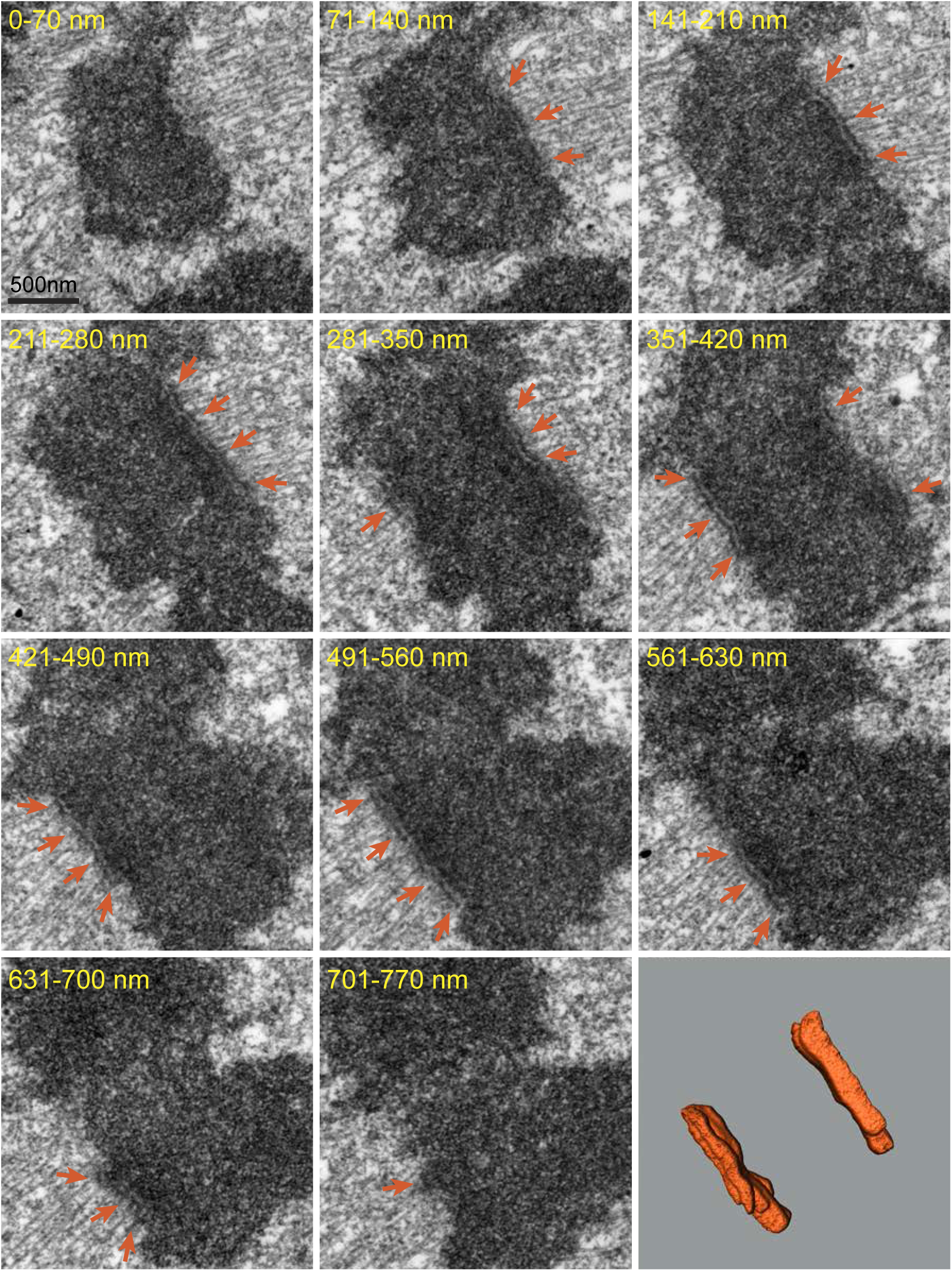
Large Kinetochore morphology in Indian muntjac metaphase. 70-nm serial EM sections from a full series through the centromere region of the chromosome. Notice the length of plate (orange arrows) and the prominent number of microtubules attached to the kinetochores. 3D reconstruction (bottom, right) of the outer layers corresponds to the kinetochores shown in EM sections.

**Figure 2 – figure supplement 3.**
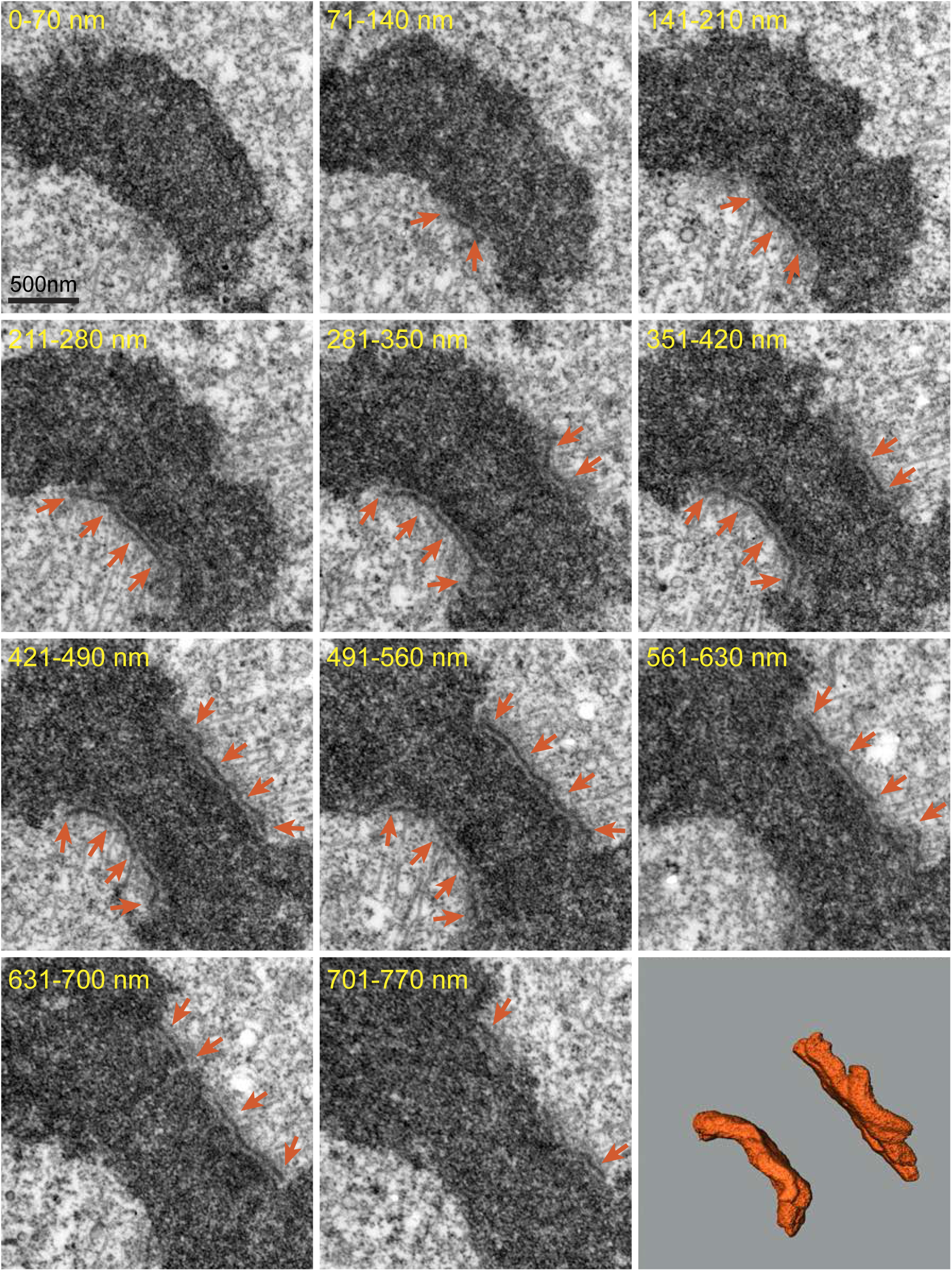
Shape and dimensions of the kinetochore remain constant in Taxol-treated IM metaphase. Serial EM sections spanning large kinetochores in Taxol-treated metaphase and 3D reconstruction (bottom, right) of the outer layers corresponds to the kinetochores shown in EM sections. No difference in shape kinetochore although the number of microtubules attached to the kinetochore plate (orange arrows) decreases after ∼15 min incubation with 10 μm-Taxol. Depth occupied by each section within the volume is shown in nanometers.

**Figure 4 – figure supplement 1.**
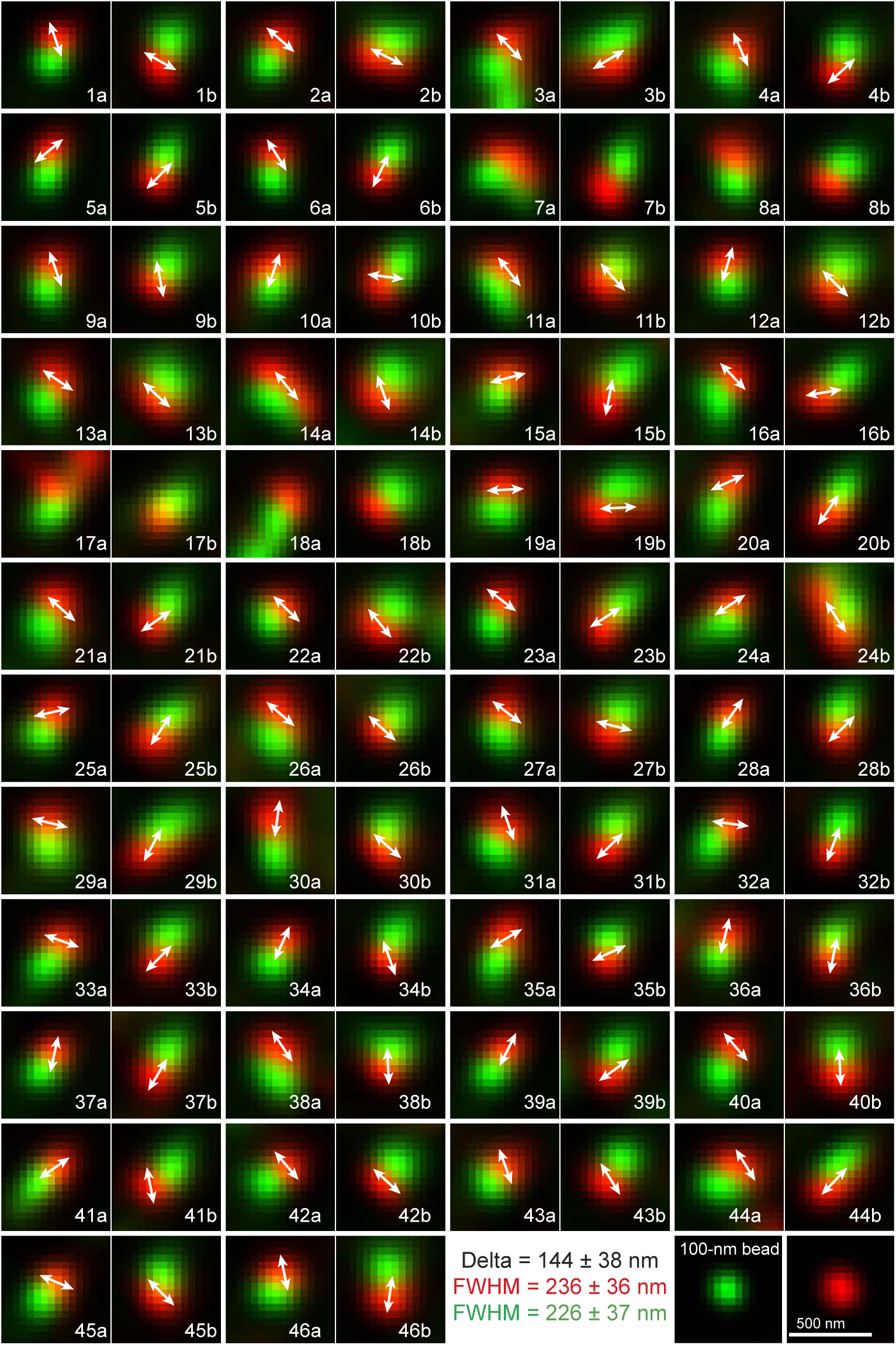
Size and shape variability of inner and outer layers of human kinetochores. 46 pairs of kinetochores in a typical Rpe1 cell during metaphase. Glutaraldehyde fixation. Numbers denote arbitrary indexes assign to chromosomes. Sister kinetochores oriented towards opposite spindle poles are marked as ‘a’ and ‘b’. Inner and outer kinetochore layers are labeled via CenpA-GFP expression (green) and immunostaining for Hec1 (red). Double arrows denote orientation of the longer axis (in XY) for Hec1 spots. Notice that kinetochore spots are larger than spots formed by 100-nm beads (shown in the bottom row). The shape of kinetochore spots and orientation of the longer axis are variable. Kinetochores on chromosomes 7, 8, 17, and 18 could not be segmented due to spatial overlap. These kinetochores were not considered in calculation of Delta, FWHM, and orientation of the larger axis. 3-D volume of the entire cell that depicts orientation of the spindle and positions of individual chromosomes/kinetochores is shown in Video 2. Kinetochore 27b is shown in Fig.4E.

**Figure 4 – figure supplement 2.**
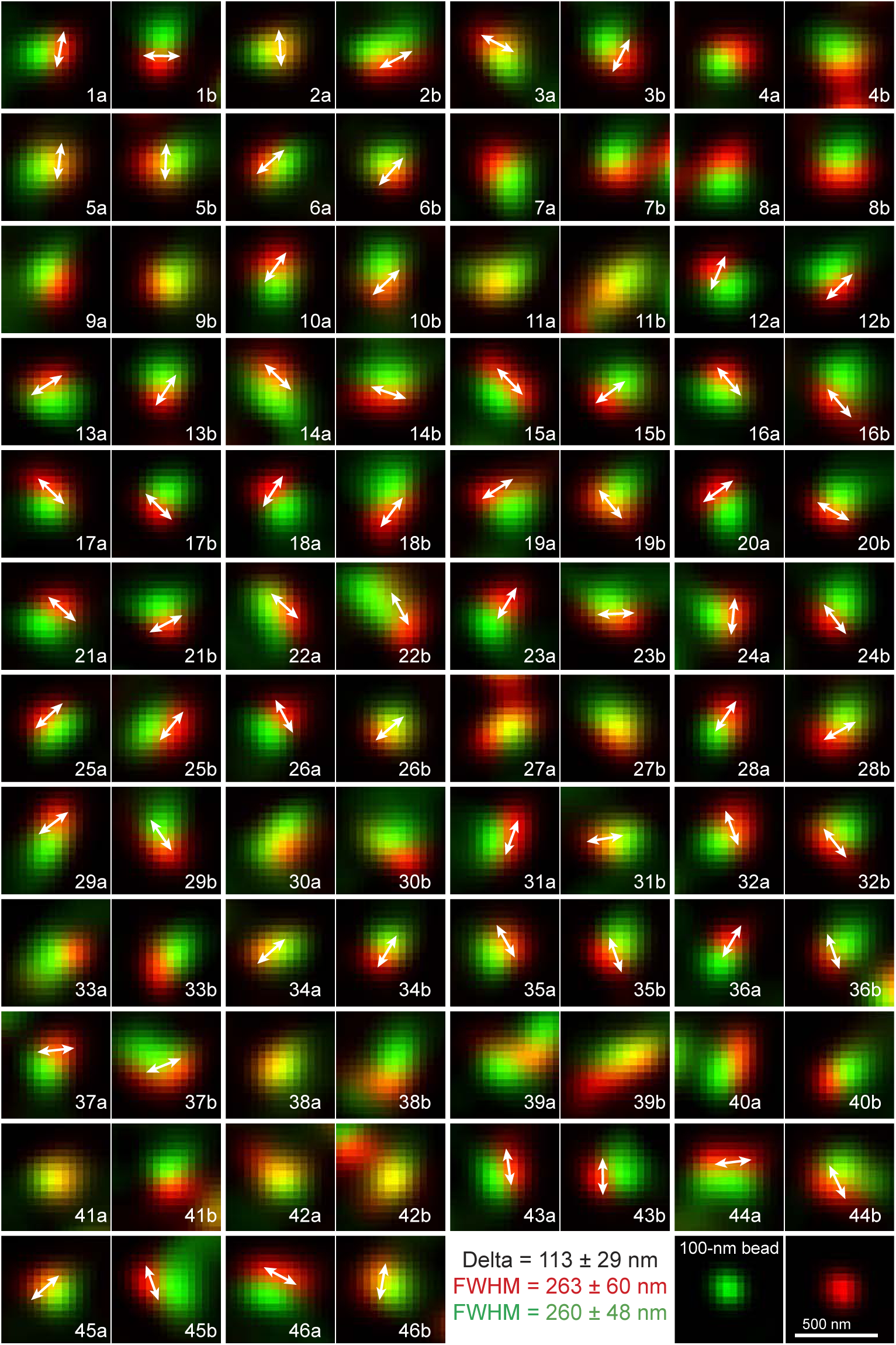
Size and shape variability of inner and outer layers of human kinetochores treated with Taxol. 46 pairs of kinetochores in a typical Rpe1 cell after 15-min exposure to 10 µM Taxol. Glutaraldehyde fixation. Numbers denote arbitrary indexes assign to chromosomes. Sister kinetochores oriented towards opposite spindle poles are marked as ‘a’ and ‘b’. Inner and outer kinetochore layers are labeled via CenpA-GFP expression (green) and immunostaining for Hec1 (red). Double arrows denote orientation of the longer axis (in XY) for Hec1 spots. Notice that kinetochore spots are larger than spots formed by 100-nm beads (shown in the bottom row). The shape of kinetochore spots and orientation of the longer axis are variable. Kinetochores on chromosomes 4, 7-9, 11, 27, 30, 33, and 38-42 could not be segmented due to spatial overlap. These kinetochores were not considered in calculation of Delta, FWHM, and orientation of the larger axis. 3-D volume of the entire cell that depicts orientation of the spindle and positions of individual chromosomes/kinetochores is shown in Video 3.

**Figure 5 – figure supplement 1.**
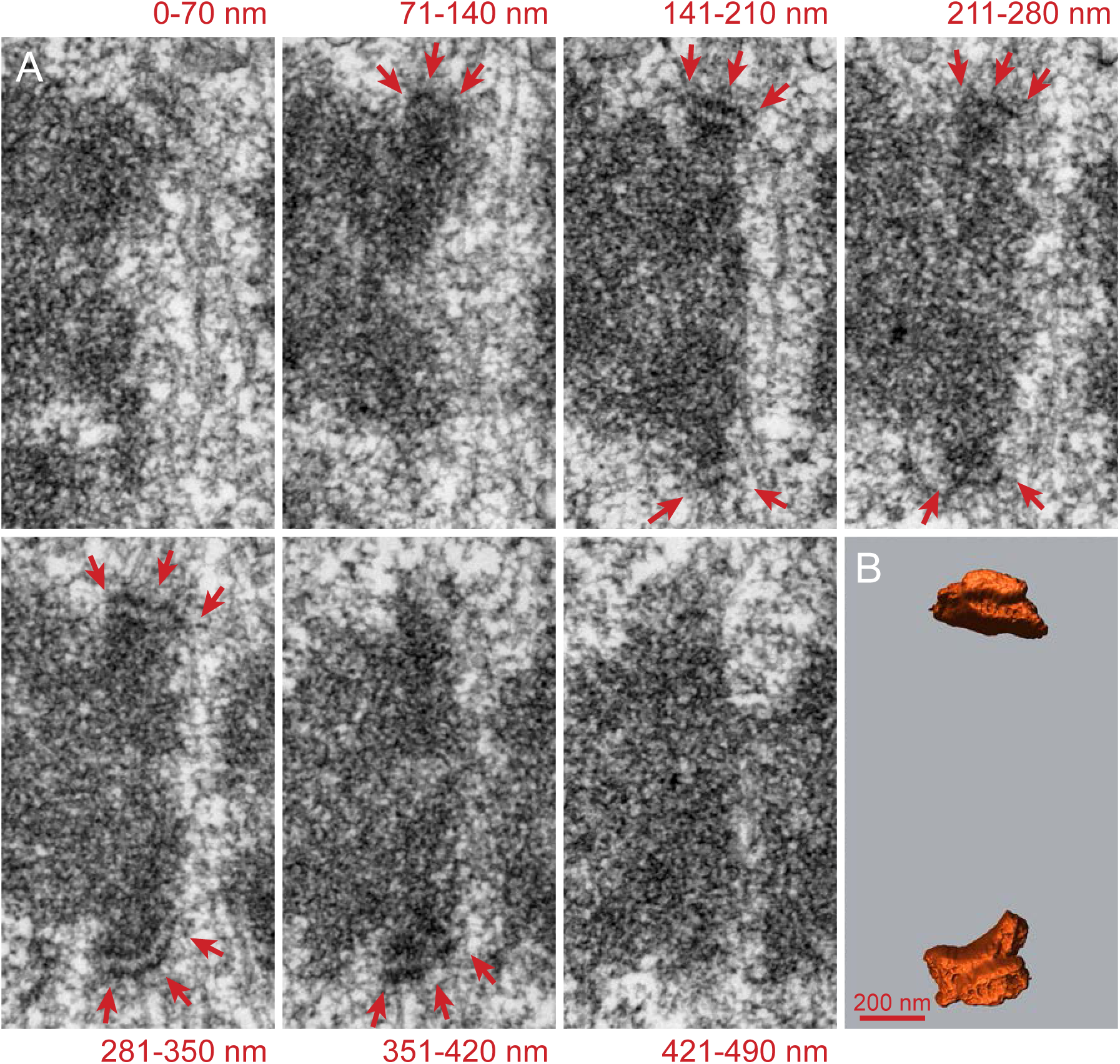
3-D reconstruction of kinetochores from a metaphase Rpe1 cell. **(A)** Selected serial 70-nm EM sections through the centromere. Numbers indicate relative depth of each section within the volume. Arrows denote the outer electron dense layers of the plates that were traced and segmented to construct a 3-D surface-rendered model shown in **(B)**. Section ‘281-350 nm’ and a different view of surface-rendered upper kinetochore are shown in Fig.5b.

**Figure 5 – figure supplement 2.**
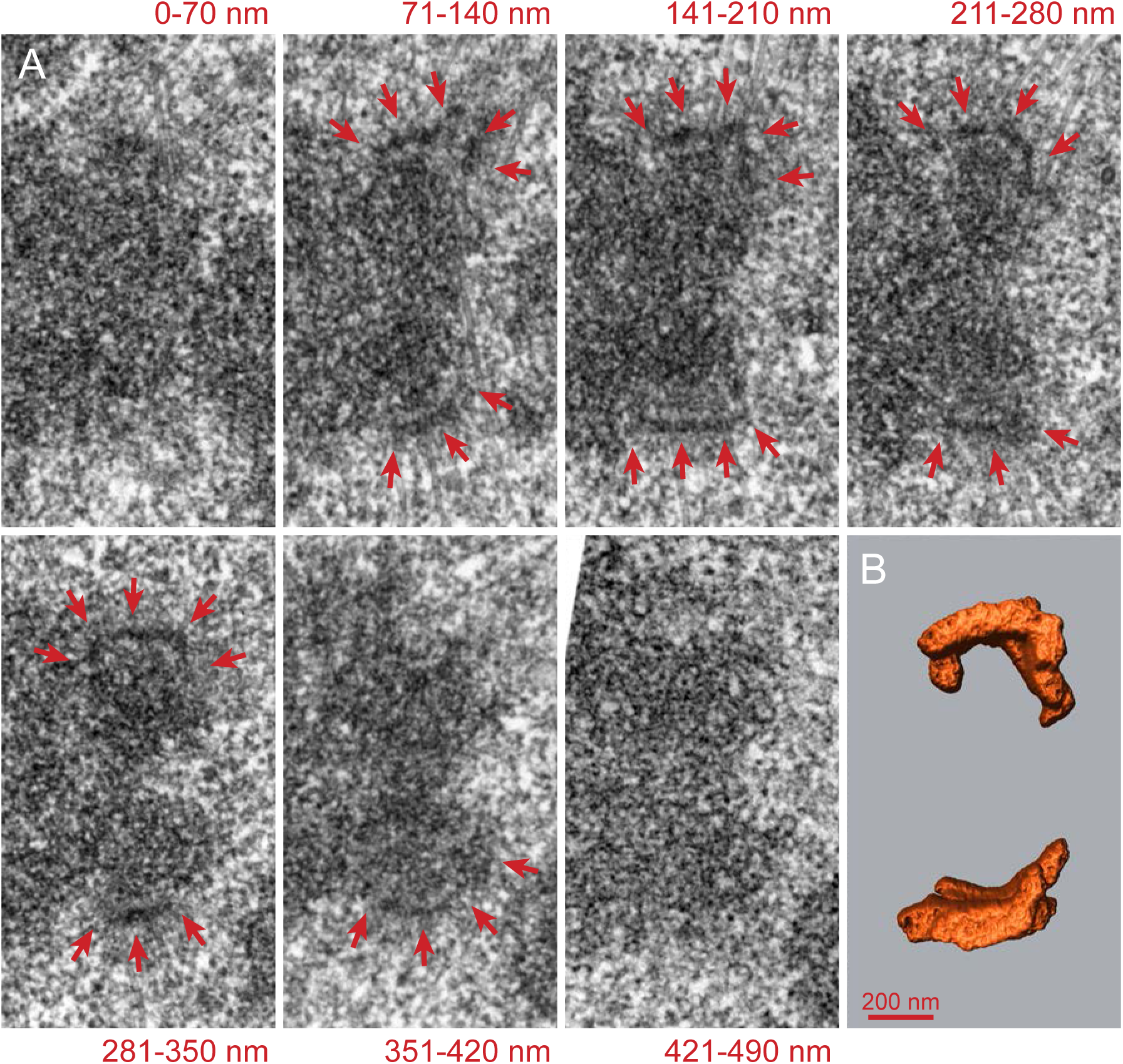
3-D reconstruction of kinetochores in Taxol-treated Rpe1 cell. **(A)** Selected serial 70-nm EM sections through the centromere. Numbers indicate relative depth of each section within the volume. Arrows denote the outer electron dense layers of the plates that were traced and segmented to construct a 3-D surface-rendered model shown in **(B)**. Section ‘211-280 nm’ and a different view of surface-rendered upper kinetochore are shown in Fig.5b.

